# Molecular Dynamics simulations of Alzheimer’s variants, R47H and R62H, in TREM2 provide evidence for structural alterations behind functional changes

**DOI:** 10.1101/536540

**Authors:** Georgina E Menzies, Rebecca Sims, Julie Williams

## Abstract

There is strong evidence supporting the association between Alzheimer’s disease (AD) and protein-coding variants, R47H and R62H in *TREM2*. The TREM2 protein is an immune receptor found in brain microglia. A structural alteration could therefore have a large effect on the protein. Crystallised structures were used as a base for both WT and mutated proteins. These subjected to 300ns of molecular dynamic simulation (MD). Results suggest structural alterations in both mutated forms of TREM2. A large change was noted in the R47H simulation in the complementarity-determining region two (CDR2) binding loop, a proposed binding sites for ligands such as APOE, a smaller change was observed in the R62H model. These differing levels of structural impact could explain the *in vitro* observed differences in TREM2-ligand binding.

**Author Summary:** A number of mutations have been found in the TREM2 protein in populations of people with Alzheimer’s and other dementias. Two of these mutations are similar in that the both cause the same coding change in the same domain of the protein. However, they both cause a very different result in terms of risk and in vitro observed changes. Why these two similar mutations are so different is largely unknown. Here we have used a in silico, simulation, approach to understanding the structural changes which occur in both of the mutations. Our results suggest that the mutation which carries a higher risk, but it less commonly observed, has a much larger impact on the protein structure than the mutation which is thought to be less damaging. This structural change is observed at a part of the protein which is thought to code for a binding loop and a change here could have a big impact on the proteins function. Further studies to investigate this binding loop could help not only a better understanding of TREM2’s role in the onset of dementia but also possibly provide a target for therapeutics.

## Introduction

The world health organisation estimate there will be 50 million dementia sufferers worldwide by 2050, with Alzheimer’s disease (AD) being the most common form [1]. Genetic studies are continually adding to the list of confirmed Alzheimer associated genes, one such study by Sims *et al* recently reported the first genome-wide evidence for the coding variant R62H in triggering receptor expressed on myeloid cells 2 (TREM2) [2]. This and other recent genetic studies have implicated the strong role of the immune system and microglial cells in the development and progression of AD [3].

The *TREM2* gene includes two genome-wide significant coding variants (R62H (Odds ratio=1.67, P=1.55×10^−14^) and R47H (Odds ratio=2.90, P=2.1×10^−12^) which are associated with an increased risk of developing AD [4–7]. Although a number of variants have been implicated in disease, they are yet to reach the level of genome-wide significance. A greater understanding of the impact of these identified variants and their impact upon the function of TREM2 in immune pathways can help with understanding how TREM2 impacts upon the development of neurodegenerative disease. TREM2 is an innate immune receptor protein which is expressed on the surface of dendritic cells, macrophages and microglia and has been shown to play an anti-inflammatory role [8]. It contains an extracellular V-type immunoglobulin (Ig) domain, a transmembrane domain which associates with the adaptor protein DAP12 for signalling and a cytoplasmic tail [9,10]. Recent studies by Zhao *et al* have shown wildtype TREM2 to bind directly to Aß with mutated forms of TREM2 showing a reduced rate of binding [11]. TREM2 has also been reported to bind to several ligands such as Apolipoprotein E (APOE) and Apolipoprotein J (APOJ) [12–14]. Subtle differences in protein secondary structure and ligand binding of R47H mutated TREM2 have previously been reported, though how these binding changes occur are not completely understood [10].

The AD associated variants are found on the surface of TREM2, this includes T66M, D87N that show suggestive association with disease and have been shown to affect the binding of ligands and other surface properties. In addition to AD, an independent number of TREM2 mutations are associated with Nasu-Hakola disease (NHD). NHD susceptibility variants (such as Q33X and Y38C [15]) are buried residues which cause complete loss of function. This segregation of disease phenotypes, table 1, when a mutation occurs within the same protein, suggests a differing effect on both the structure and function of TREM2 [10].

**Table 1.** Known variants of TREM2 and the suggested impact [10,16]

| TREM2 Variant | Surface Expression Changes | Ligand Binding Changes | Signalling Response | % solvent Accessibility | Implicated in |
| --- | --- | --- | --- | --- | --- |
| Y38C | Reduced | Ablated | N/A | 7.6 | NHD |
| R47H | None | Reduced | Reduced | 32.5 | AD |
| R62H | None | Reduced | Reduced | 48.0 | AD |
| T66M | Reduced | Ablated | None | 0.0 | NHD |
| N68K |  |  |  | 77.2 | AD |
| D87N | None | Reduced | Increased | 66.7 | AD |
| **T96K** | Reduced | Increased | Increased | 26.7 | AD |
| **V126G** |  |  |  | 0.0 | NHD |

In order to investigate the structural impact of the R47H and R62H mutations and predict possible loss-of-function we carried out an *in silico* study of the binding domain of the protein containing the mutations. Here we describe the results of this study and in particular the similarities and differences between the two models. Results suggest a greater effect on the binding loops by the R47H mutation, fitting with previous studies [10].

## Results

The wildtype TREM2 protein’s extracellular ligand binding domain (ECD) contains amino acids 19-134 which have been crystallised from a mammalian cell system [10]. The rest of TREM2 contains a signalling peptide (1-18), a membrane spanning region (175-195) and a cytosolic tail (196-230) [17]. These all form important roles in the protein but as they are not suggested to be used in the binding process, the process which is affected by the AD risk mutations, and they are not crystallised, they have been excluded from this study. The domain under investigation is a V-type Ig domain which contains nine B-strands and two short a-helices, all of which are characteristic of an Ig protein domain. Both mutations, R47H and R62H, can be found on the protein surface, which is suggested to be how they affect TREM2’s binding abilities, in particular its ability to bind to APOE and APOJ [12,13]. The Have your Protein Explained server (HoPE) was used to investigate the possible mutational effects prior to any simulations being run [18]. Results from the server suggest that the wildtype amino acid (arginine) at position 47 forms a hydrogen bond with amino acids at positions 66 (threonine) and 67 (histidine) which would not be possible with the histidine mutation in this position. These bonds may be important for protein structural integrity. The wildtype residue is conserved at position 47, though histidine is observed here in some species. Residue 62 on the other hand is less well conserved, but histidine is not observed here. There is an obvious loss of charge and size with mutated R62H, shown schematically in figure 1. The SIFT online tool was used to predict the tolerance of the two mutations in the protein, this does not predict the effect of binding, or function, but whether the mutation will be tolerated in the protein structure. Results from this show the R47H mutation to be tolerated with a score of 0.06 and the R62H mutation to be tolerated with a score of 0.10, this was based on 13 sequences. A score of <0.05 would result in a damaging prediction [19]. The I-mutant server results showed a decrease in stability for both the R47H and R62H mutations [20].

**Figure 1.**
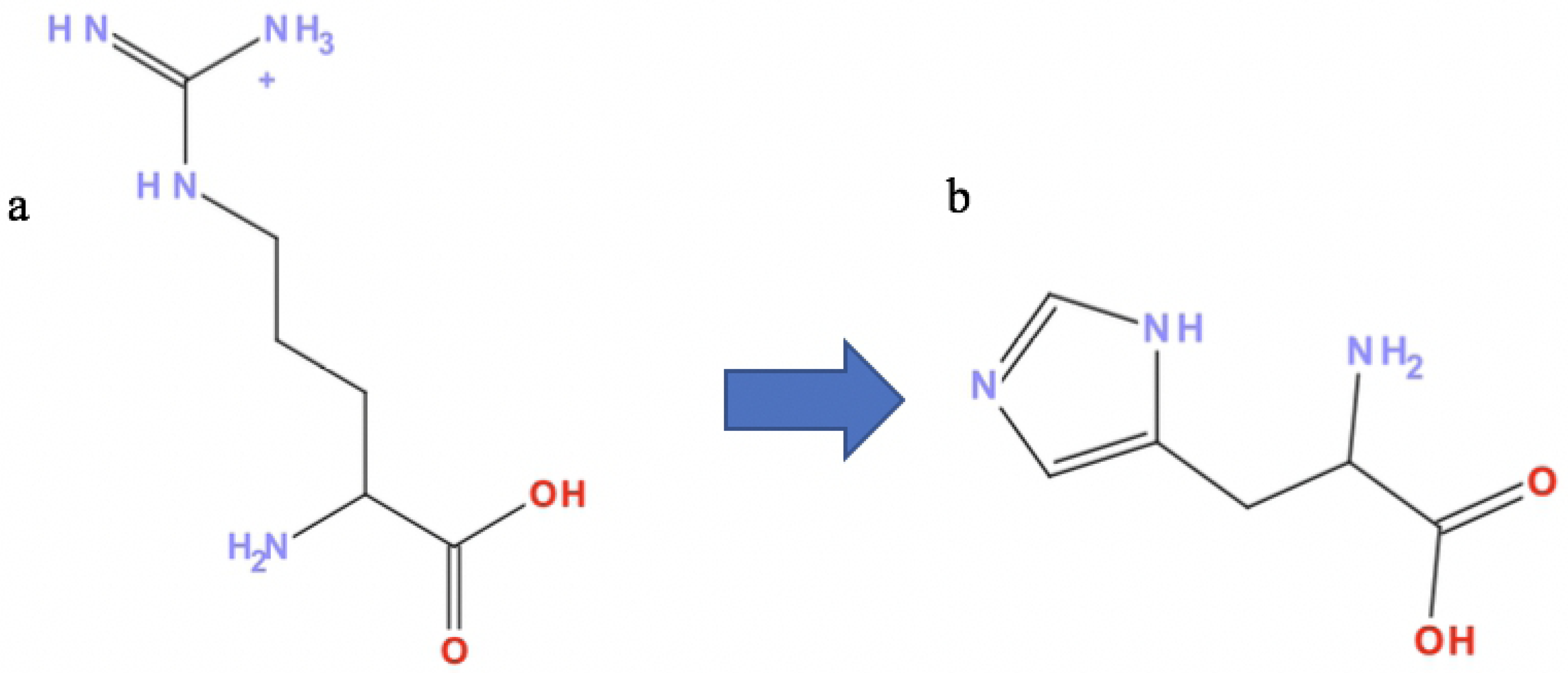
Schematics of the (a) wildtype and (b) mutated amino acids

MD simulations were run, in triplicate, for the WT and mutated proteins, the stability of the simulations were checked, and volume, pressure and root mean square deviation (RMSD) remained stable throughout thus giving confidence in the model systems. Figure 2 depicts the structure of TREM2 which has been modelled and run through the MD simulations. The complementarity-determining region (CDR) loops, which are suggested to be key for the ligand binding process [21,22], are coloured as follows; red for the CDR1, green for CDR2 and purple for the CDR3 loop. The two mutated sites are shown in dark blue, both are close to the CDR1 and CDR2 loops, with R47H actually being found in the CDR1 loop.

**Figure 2.**
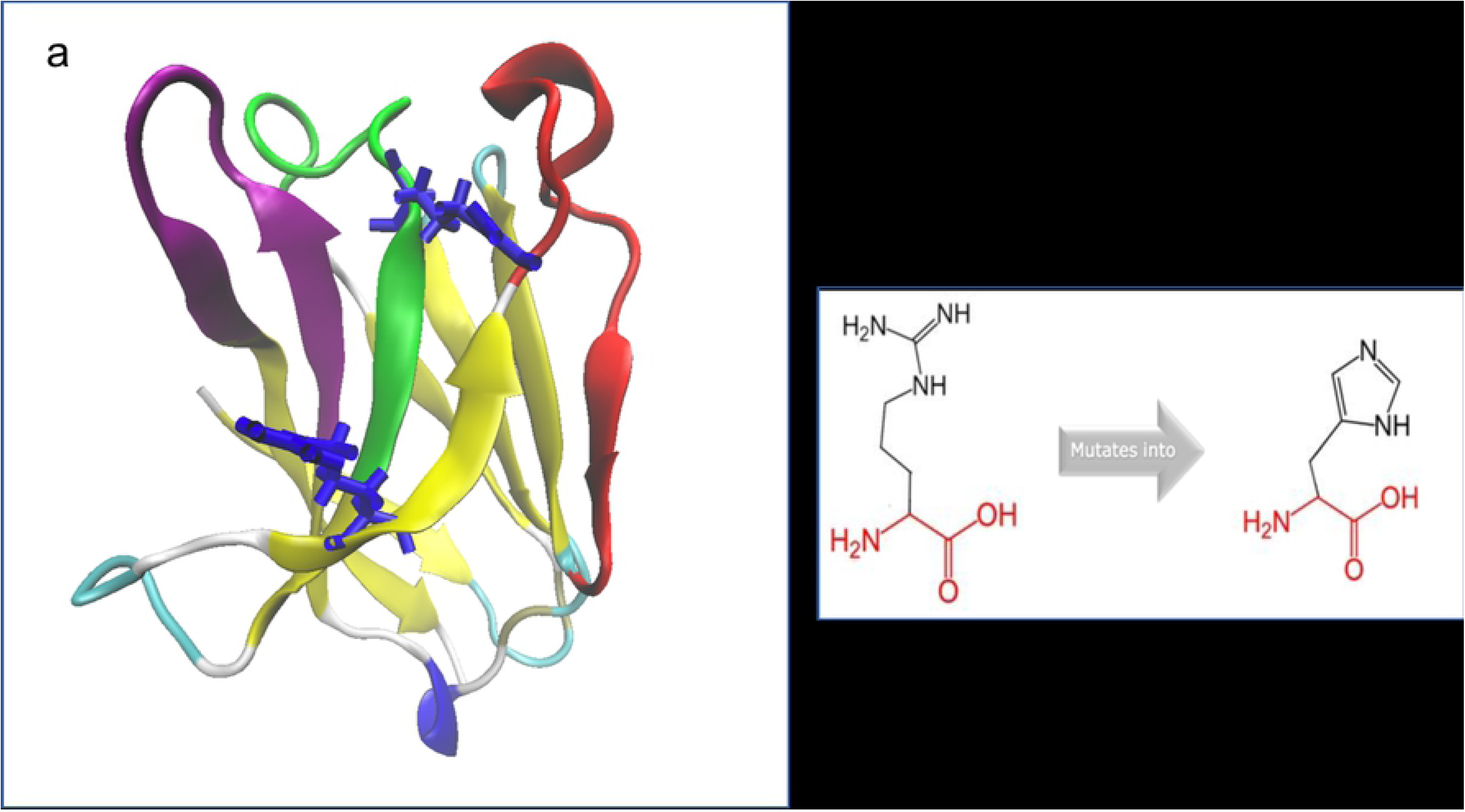
Wild type TREM2, the structure is depicted as a cartoon style with secondary structure colouring. CDR loops are coloured as follows; CDR1 = red, CDR2 = green, CDR3 = purple, the position of the two mutated sites are coloured in dark blue and shown in full.

Mutations could be impacting the local or the global structure of TREM2. Local structural changes were first investigated in the three molecular models. The region surrounding both mutations, amino acids 43-65 were viewed, figure 3. The R62H mutation alters this local structure with a shift in the beta sheet and a large movement of the random coil. The R47H mutation does not appear to alter this local structure in any way. As well as altering the local structure the flexibility of the individual residue, i.e. the amount of movement it has, was also altered for the R62H mutation. Results show the WT and R47H have a flexibility of 0.23 +/- 0.02 and 0.01 respectively at amino acid 62, the R62H mutation on the other hand has a reduced flexibility of 0.17 +/- 0.01. There is also, to a lesser extent, a reduction of flexibility across neighbouring amino acids which surround the R62H mutation.

**Figure 3.**
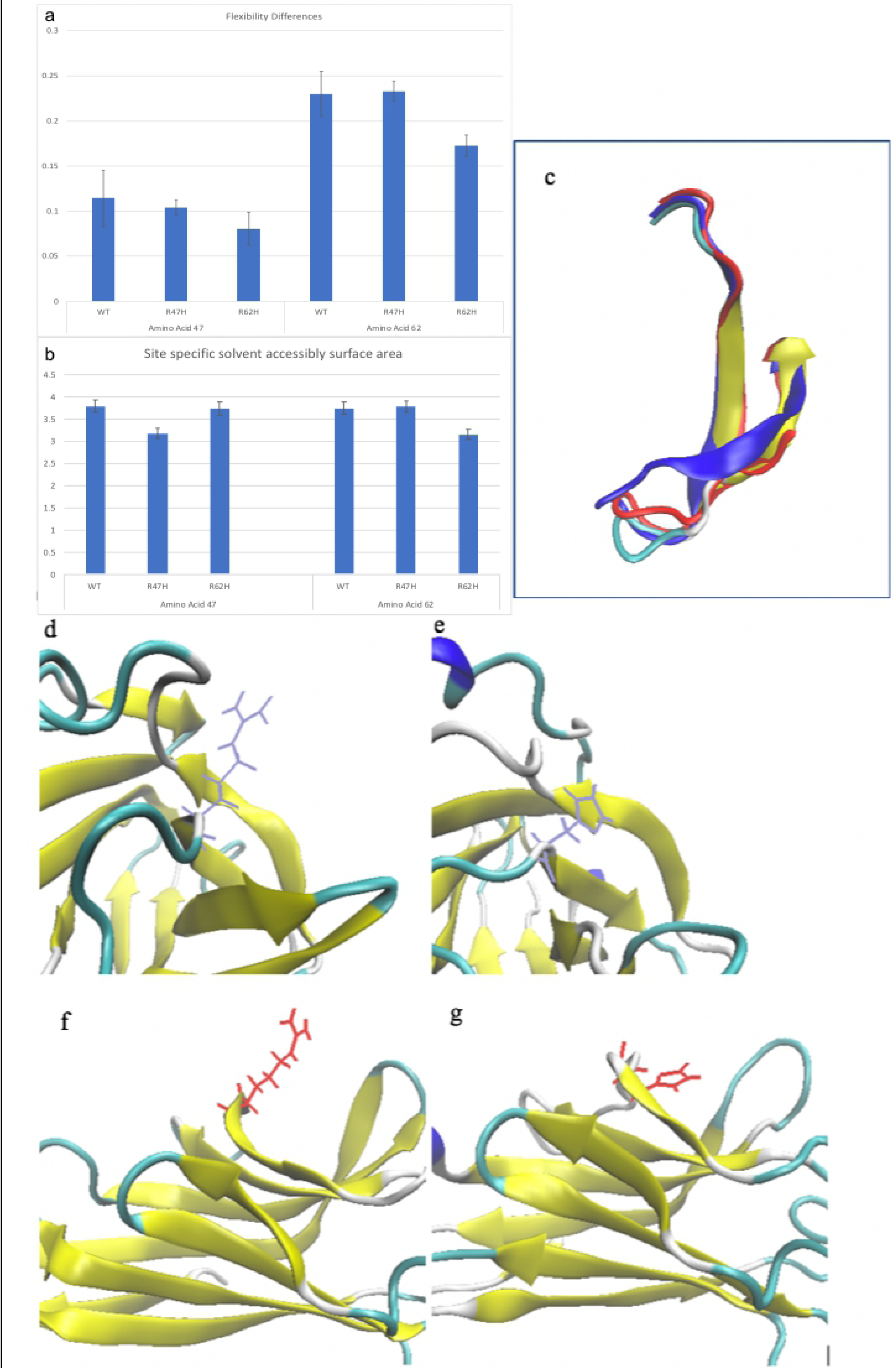
Graphs of the flexibility changes for the wildtype and mutated proteins at both sites as well as the point specific SASA are shown in a and b. c depicts the local level structural alteration with the type in secondary structure colours, R47H in red and R62H in blue. d-g show the wildtype and mutated acids positioning for the R47H wildtype, mutation, and the R62H wildtype and mutation respectively.

Arginine, which is present in the wildtype protein at both positions, is a long and stretching amino acid with a chain of carbons and nitrogens. Histidine, which is the mutated form of both variants, is a ring strucutre, with less avaliblity for hydrogen bonding. MD simulation results show a change in positioning of the wildtype to mutated amino acid, the wildtype pretuding from the molecule in both cases and the mutated amino acid being visually far more buried within the structure, figure 3 (d-g).

Solvent accessible surface area (SASA) for the whole protein, and the individual mutated residues were measured. Overall SASA was reduced from 71 to 70, this small change is not significant and may not have any effect on the protein function. Amino acid specific SASA was measured for the WT and mutated proteins, at the 47 and 62 sites. Here a SASA change can be seen at the mutated site in each protein, with a reduction of SASA, figure 3b.

A final measurement of the distance between the two mutation sites (taken to show structural shrinkage in the protein) was measured. Again, a reduction was seen here in the R47H and R62H mutations when compared to the WT.

Significant structural alteration can be seen in the CDR2 loop, figure 4 shows the R47H mutation to cause a loss of beta sheet and a changing of alpha helix position in this binding loop. The effect of the R62H mutation is subtler, where there is a change in loop structure to the left and right of the alpha helix. There is a further effect on the flexibility of the loop, figure 4c, here the mutations differ with the R47H becoming more flexible and the R62H mutation less flexible, when compared to the wild type simulation. Significant structural alteration can be seen in the CDR2 loop, figure 4 which contains the results of all three simulations shows the R47H mutation to cause a loss of beta sheet and a changing of alpha helix position in this binding loop. The effect of the R62H mutation is subtler, where there is a change in loop structure to the left and right of the alpha helix. All three images show the loop in the same position and therefore should all be identical if no structural change was observed.

**Figure 4.**
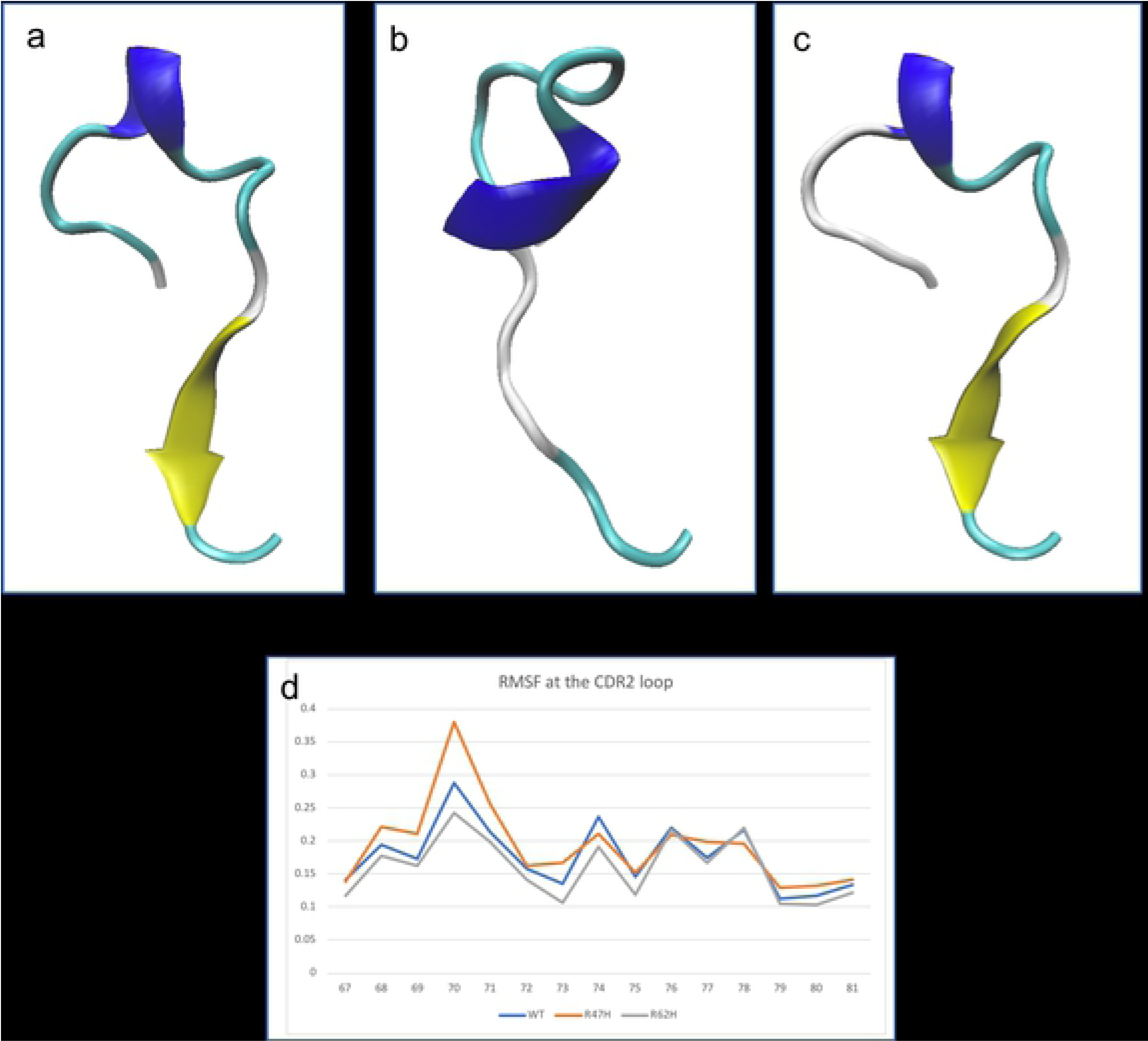
The TREM2 CDR2 loop which spans amino acids 67 to 81, (a) wild type, (b) R47H mutation, (c) R62H mutation (d) the RMSF – or flexibility of the residues in the CDR2 loop.

## Discussion

In this study we present structural findings and subsequent functional predictions from two AD associated genome-wide significant mutations in TREM2. The R62H mutation which is more common in the Caucasian population with OR=1.67; P=1.55×10^−14^ [2] whereas the rarer mutation, R47H, carries a far greater risk (OR= 2.90; P=2.1×10^−12^ [4]). Both mutations are found within exon 2, the region which is predicted to encode for the ligand-binding domain, and both are missense mutations causing a coding change from the wild type arginine to a histidine. Previous studies have shown that disruptions to the protein in exon 2 are likely to cause TREM2 signalling problems or a loss or a decrease in protein function. The functional impact of both variants has been discussed at an *in vitro* and *in vivo* level, but here we present a study which aims to identify the structural cause behind the functional alterations. Previous studies have identified R47H as having a greater functional effect than R62H even though the same mutation is observed and they are in very close proximity to each other, why this difference happens is largely unknown [10]. Other TREM2 variants have been suggested to have an effect in AD, but so far only these two coding variants have reached genome-wide significance and so are the only two studied here [10,23]. Previous studies have looked into the structure of TREM2, one such study, published in 2014, by Abduljaleel *et al* performed MD simulations on the R47H variant. Their simulations ran for just 10ns and they presented results which suggested a possible alteration to binding loops and overall stability alteration. This short simulation time may not have been long enough to view any large or distal impact and this study builds upon those results and expands their hypotheses [24].

The differing results from SIFT and I-Mutant suggest an overall tolerance of the mutation, but a local structural shift. The overall tolerance of the mutations is key, the proteins remaining stable, and tolerating the mutation means the mutated protein is likely to perform a partial function as is predicted *in vitro*. The protein is also likely to be expressed on the cell surface still, as is also predicted. This is supported by the simulation results which suggest no change in the global protein flexibility or SASA. The local structure shows a greater change, beginning with the positioning of the amino acid. The wildtype for both mutations are found outstretched from the proteins binding domain, here they could perform key functions in binding, it has been suggested that the positive amino acids such as these play an important role [25]. The mutated residues are neutral in charge, provide less opportunity for hydrogen bonding and are buried within the binding domain, this alone causes an impact on TREM2’s ability to bind to ligands such as APOE. Further to this local change, both mutations are found in the vicinity of the binding loops of CDR1 and CDR2, R47H lies on CDR1 and R62H between the two loops. These, and other putative AD mutations, are found on the surface of the protein where they may affect TREM2’s ability to bind and function.

Solvent-accessible surface area, SASA, is important when considering rates of reactions which require a protein-protein or protein-ligand interaction and so a change in the SASA of either of these two amino acids which could be key in the binding process should be considered a detrimental effect and results showed a reduction in SASA at the mutated residue for both models [26]. A further result of note is the reduction the distance between the amino acids for the R62H model, this measurement suggests a reduction in overall protein size and a loss of shape, two things which are again key for function. Sudom *et al* recently published a paper which showed mutated R47H protein to contain a remodelled helix in the CDR2 loop, though their crystal structure is missing residues 76-81[17]. This study supports an altered helix structure in the CDR2 loop, we also see a loss of the beta sheet structure which is replaced by a random coil. A random coil is far more variable and could explain why they were unable to resolve this region of the protein and the crystal structure is missing this region. This TREM2 domain also contains three possible N-glycosylation sites, one of which is at position 79, the alteration in structure here could be effecting the ability of TREM2 to undergo translational modification and could explain the altered glycosylation seen in vitro in the R47H mutated form [27].

Park *et al* recently showed that the R47H mutation in TREM2 resulted in a decreased protein stability, based on our models this may due to the large alteration in the CDR2 loop structure [27]. Another study by Atagi *et al* presented strong evidence for the binding of TREM2 to APOE, and more interestingly a lack of binding when the R47H mutation was present [14], this is further supported by Yeh *et al* who measured a decrease in TREM2’s ability to bind CLU/APOJ and APOE when the R47H and R62H mutations were present. Their results support our difference in binding loop loss between the two mutations as they observed less of a decrease in binding with the R62H mutation [12]. This binding loop degradation we observed may be the key to understanding the functional effect these mutations are having on the protein.

The evidence shown here correlates with previous studies which indicate a binding change when the R47H mutation is present. We present novel findings which show the R62H mutation to have a structural effect on the same region of the protein albeit to a lesser extent. This provides insight and support to the studies which show less of a decrease in binding ability with the R62H mutated protein compared to the R47H mutated form. Understanding the structural and functional changes which occur in this AD associated protein increase our knowledge of the mechanisms behind the processes which cause AD and as a result provide more novel drug and therapeutic targets.

## Materials and Methods

The immunoglobulin domain for the TREM2 protein has previously been crystallised [10], both mutations were added to the structure using the modify protein function in the Accelrys software, Discovery studio. The wildtype protein (WT) and the two mutated structures were subjected to over 300ns of molecular dynamics (MD) simulations. MD was carried out using the GROMACS [28] software suit using the Amber03 [29] in built force field parameters. All protein structures were placed in a cubic box, solvated using TIP3P water molecules and neutralised using Cl^−^ ions. The particle mesh ewald (PME) method was used to treat long-range electrostatic interactions and a 1.4 nm cut-off was applied to Lennard–Jones interactions. All of the simulations were carried out in the NPT ensemble, with periodic boundary conditions and at a temperature of 310K. There were three-steps to each simulation. 1; Energy minimisation, using the steepest decent method and a tolerance of 1000KJ^−1^ nm^−1^. 2; Warm up stage of 25 000 steps at 0.002ps steps, during this stage atoms were restrained to allow the model to settle. 3; Finally, a MD stage run for a total of 300ns. Root mean square deviation (RMSD) was monitored along with the total energy, pressure and volume of the simulation to check for stability.

Resulting structures were analysed for flexibility using the gmx rmsf and hydrogen bonding using gmx hbond (both available within the GROMACS suite) all proteins were visualised for structural differences using VMD. Further to this prediction of the functional effect and stability analysis was carried out using three online servers, HoPE, SITF and I-mutatnt [18–20,30]. HoPE analyses the impact of a mutation, taking into account structural impact, and contact such as possible hydrogen bonding and ionic interactions. The SIFT software predicts tolerated and deleterious SNPs and identifies any impact of amino acid substitution on protein function and lastly, I-Mutant is a neural-network based prediction of protein stability changes.

Statistical normality in distributions such a rmsd, energy, pressure, volume etc, were tested for using the Anderson-Darling test. All were not normally distributed and so all statistical differences between the wildtype and mutated simulations were calculated using the Mann-Whitney *U* test.

## Acknowledgements

Part-funded by the European Regional Development Fund through the Welsh Government

